# An *in vitro* regeneration system with efficient rooting in sweet orange (*Citrus sinensis*) supports recovery of transgenic plants

**DOI:** 10.64898/2026.06.16.732047

**Authors:** Juel Datta, Sudipta Das Bhowmik, Brett Williams, Stephanie C. Kerr

## Abstract

*In vitro* regeneration of *Citrus* plants is a widely used method, however, induction of adventitious roots from regenerated shoots remains a major bottleneck, limiting the recovery of healthy plants for commercial production and genomic research for crop improvement. We established an in vitro regeneration system producing profuse, healthy roots for sweet orange (*Citrus sinensis* cv. Benyenda) by optimising combinations and concentrations of auxins. Prior to optimising the rooting media (RTMs), we obtained a shoot regeneration rate of 90.6% from sweet orange epicotyl explants using a cytokinin, 6-benzylaminopurine (BAP). Across twelve auxin-supplemented RTMs containing different concentrations of indole-3-butyric acid (IBA) and/or 1-naphthaleneacetic acid (NAA), rooting percentages ranged from 8 - 87.5%. The combination of IBA 1.0 mg L^-1^ and NAA 0.1 mg L^-1^ promoted the best overall performance, 75 ± 7.2% rooting percentage with healthy, callus-free roots (≥5 cm in length), whereas other RTMs with other auxin combinations induced callus and limited root elongation. The best-performing SRM and RTM were subsequently used for selection and recovery of transgenic sweet orange lines carrying an empty CRISPR/Cas9 construct, resulting in an 4.8% transformation efficiency. Both transgenic and non-transgenic rooted plantlets were successfully acclimatised under glasshouse conditions with a survival rate of 90%. This enhanced regeneration system overcomes rooting bottleneck and improves plant survival,enabling faster recovery of transgenic citrus lines within four months. It supports accelerated development for commercial applications and advances in citrus genetic improvement.

## Introduction

Citrus crops are among the most economically important fruit tree species worldwide, encompassing oranges, mandarins, lemons, grapefruits and limes. They are in high global demand and are widely valued for their use in the fresh fruit and pharmaceutical industries ^[1]^. Citrus products consumed worldwide are predominantly derived from commercial citrus varieties, with sweet orange (*Citrus sinensis*) being the dominant species cultivated ^[2–4]^.

*In vitro* regeneration is an efficient approach commonly used for large-scale production of elite citrus varieties and to facilitate trait improvement via genetic transformation in breeding programs ^[5–7]^. The success of plant regeneration depends on many factors, including explant types, tissue culture conditions, and media composition, of which the composition of plant growth regulators (PGRs) in the media is most important ^[8–11]^. Juvenile epicotyl explants are widely used in citrus regeneration and transformation methods due to their high morphogenic competence, rapid shoot regeneration capacity and reduced somaclonal variation-*Citrus* species facilitates the propagation of plant from seed-derived tissues that are true-to-type to the mother plant ^[12–16]^. Optimal combinations of PGRs are required for efficient citrus regeneration and the recovery of transgenic lines. Although many studies have focused on optimising culture media to overcome the recalcitrant nature of citrus and its genotype-specific responses to *in vitro* conditions, regeneration efficiency remains inconsistent due to differences in tissue responsiveness to PGRs ^[11]^.

*In vitro* rooting from regenerated shoots is one of the most crucial stages in the regeneration system of many perennial fruit trees including citrus ^[17]^. The success of an *in vitro* regeneration system is ultimately determined by the survival of plantlets during acclimatisation. In citrus, inefficient rooting of regenerated shoots is considered a major bottleneck that prolongs tissue culture timelines, limits the recovery of vigorous plants, and often necessitates alternatives such as micrografting ^[18–20]^, which are labour-intensive, time-consuming, and dependent on rootstock availability and compatibility ^[21,22]^. Root induction, subsequent root growth, and overall plantlet vigour are highly sensitive to the type and concentrations of PGRs ^[23,24]^. As a result, regenerated shoots may fail to root or root only at low frequencies;for instance, transgenic ‘Duncan’ and ‘Hamlin’ cultivars failed to root in media that supported rooting in ‘Carrizo’ citrange and Mexican limes were successfully rooted ^[25]^. These genotype- and protocol-dependent responses of citrus species highlights that no single tissue-culture method is universally applicable. Therefore, optimising PGR regimes and rooting media composition is essential to achieve consistent rooting and reliable acclimatisation, enabling large-scale recovery of plants, including genetically modified lines, while reducing dependency on grafting approaches.

The use of regeneration systems in biotechnological approaches, such as CRISPR-based genome editing, facilitates trait modification ^[26–28]^. The successful recovery of CRISPR-edited plants in Citrus remains largely dependent on *Agrobacterium*-mediated transformation systems that require an efficient *in vitro* regeneration method ^[5,29–32]^. Currently, low shoot regeneration along with poor rooting limits the overall plant survival rate, which restricts the broader application of *Agrobacterium*-mediated transformation protocols ^[13,17,33]^.

Although varying combinations of PGRs have been tested for *in vitro* citrus regeneration, very few studies have specifically focused on optimising rooting responses to different PGRs. Considering the ongoing challenges associated with citrus regeneration and the recovery of transgenic plants, we aimed to improve rooting efficiency from regenerated shoots in a commercial sweet orange cultivar. The best-performing media were further trialled for the successful recovery of transgenic citrus plants.

## Materials and methods

### Plant materials and seed germination

Mature seeds were harvested from the fruit of the commercial citrus species, sweet orange, *Citrus sinensis* cv. ‘Benyenda’. Seeds were thoroughly washed with sterilised distilled water (SDW) to remove any pulp and sterilised with 2% v/v bleach (NaOCl) by soaking for 10 minutes, then rinsed with SDW three times and air dried. The outer seed coats were removed manually inside an laminar flow chamber and the seeds were then aseptically cultured on Murashige and Tucker (MT) medium ^[34]^ supplemented with 3% (w/v) sucrose and 0.8% (w/v) agar and incubated at 25 ± 1°C for 2-3 weeks in the dark followed by 2 weeks under a 16-h photoperiod. Based on a pre-trial in our lab facilities, an optimal light condition using cool-white LED tubes (Valoya) with an intensity ranging from 100 to 200 µmol s^-1^ was used to maximise seedling growth, development and subsequent regeneration. Seedlings approximately 5-6 cm long were cut vertically to excise 1 cm long epicotyl explants under aseptic conditions.

### Shoot regeneration: media and culture conditions

Excised epicotyl explants were cultured on MT basal medium containing 3% (w/v) sucrose and solidified with 0.8% (w/v) agar. Different combinations and concentrations of the cytokinin 6-benzylaminopurine (BAP) and the auxins indole-3-acetic acid (IAA) and 2,4-dichlorophenoxyacetic acid (2,4-D), were added to create different shoot regeneration media (SRMs). All media were adjusted to pH 5.8 prior to autoclaving, and filter-sterilised PGRs were added to cooled, sterile medium to prepare following SRMs: SRM1 – BAP 2.0 mg L^-1^, SRM2 – BAP 3.0 mg L^-1^, SRM3 –BAP 2.0 mg L^-1^ + IAA 0.5 mg L^-1^, and SRM4 – BAP 2.0 mg L^-1^ + 2,4-D 0.25 mg L^-1^. Explants were placed on SRMs for shoot initiation and sub-cultured onto fresh medium of the same composition every 2 weeks for up to 6 weeks before transfer to rooting media (RTMs). Each of these SRMs (treatment) was comprised of a minimum of three replicates, each with at least 32 epicotyl explants. Cultures were maintained at 25 ± 1°C under a 16-h photoperiod with cool white LED lighting (100-200 µmol s^-1^) (Valoya). The shoot regeneration rate (%) was calculated as the following formula:

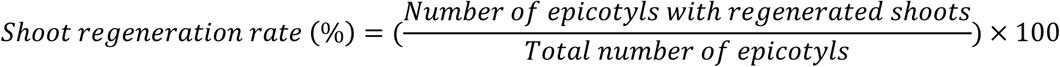

### Rooting of regenerated shoots

Rooting was induced by subculturing regenerated shoots onto MT-based RTMs prepared at either half-strength or full-strength with 1.5% or 3.0% (w/v) sucrose, respectively. RTMs were supplemented with the auxins indole-3-butyric acid (IBA) and 1-naphthalene acetic acid (NAA) at various concentrations and combinations. A full factorial design was used with IBA at 0, 0.5, 1.0, and 2.0 mg L^-1^ and NAA at 0, 0.1, and 0.5 mg L^-1^, yielding twelve RTMs in total. Regenerated shoots of around 1–2 cm long were excised aseptically and transferred to RTMs. Each RTM (treatment) comprised three replicates, each with 8 shoots. Data on root number, root length, and root morphology were collected weekly between 2 and 8 weeks after transfer. Culture conditions were held constant at 25 ± 1°C with a 16-h photoperiod. The rooting efficiency in terms of rooting percentage was determined by dividing the total number of regenerated shoots inducing roots with the total number of regenerated shoots used for rooting, multiplied by 100.

### Acclimatisation of plantlets

Regenerated plantlets with healthy roots (>3 cm in length) were transferred to plastic pots containing an autoclaved potting mixture supplemented with Osmocote (10:1 ratio). Plantlets were rinsed thoroughly with SDW to remove residual culture medium prior to transplanting. The potted plantlets were covered with transparent plastic sheets to maintain high humidity for 2 weeks. After acclimatisation was visually confirmed the plastic covers were removed. During the acclimatisation process the plants were grown in a controlled environmental condition at a constant temperature of 25 ± 1°C with a 16-h photoperiod. Survival of the plantlets was evaluated after 4 weeks.

### *Agrobacterium*-mediated epicotyl transformation and regeneration

A modified pCAMBIA2300 binary vector, renamed pS203 (**Supplementary Fig. S1;** modified and provided by Sally Roden, Waterhouse group, Queensland University of Technology), was used in this study. The vector contains a Cas9 expression cassette without incorporated guide RNAs (gRNAs). Cas9 expression is driven by the CaMV35S promoter and terminated by the NOS terminator. This binary vector functions as an empty CRISPR/Cas9 construct and confers kanamycin resistance for the transformation of sweet orange epicotyl explants without modifying the plant genome. The use of this vector will facilitate the establishment of a CRISPR-mediated gene delivery system in sweet orange, employing the tRNA–gRNA array architecture reported by Xie et al ^[35]^. Plasmids containing the empty CRISPR/Cas9 construct were introduced into *Agrobacterium tumefaciens* strain GV3101 by electroporation using a Gene Pulser^TM^ (BIO-RAD) following 1.80 V, 200 Ω, 24 µFD settings.

In this study, we used an *Agrobacterium*-mediated epicotyl transformation method according to Jia et al ^[36]^ with minor modifications to deliver an empty CRISPR/Cas9 construct. Briefly, *Agrobacterium* harbouring the empty CRISPR/Cas9 construct were recovered and grown overnight at 28°C in 5 mL of LB media containing kanamycin (50 mg L^-1^) and rifampicin (100 mg L^-1^). The next day the culture was centrifuged and diluted in 20 mL of co-cultivation medium (CM; MT medium supplemented with 3.0 mg L^-1^ BAP, 0.25 mg L^-1^ 2,4-D, 100 µM acetosyringone, and 3.0% (w/v) sucrose) and the OD_600_ value of the *Agrobacterium* suspension was adjusted to 0.6. The excised sweet orange epicotyl explants were suspended in CM, shaken gently at 50 rpm for 2 hours and then resuspended in *Agrobacterium* suspension for 20 minutes. The epicotyls were transferred onto sterile filter paper in a new Petri dish and dried for 3-5 minutes, then transferred onto solidified CM (solidified with 0.8% (w/v) agar). After *Agrobacterium* co-cultivation, explants were incubated in darkness for 72 hours at 25 ± 1°C, then transferred to SRM2 selection medium and maintained under a 16-h photoperiod at 25 ± 1°C. SRM2 was supplemented with kanamycin (100 mg L^-1^) and cefotaxime (250 mg L^-1^) to suppress bacterial overgrowth; this modified medium was further referred as SRM2+. Explants were sub-cultured to fresh SRM2+ every 2 weeks with the kanamycin concentration gradually reduced to 50 mg L^-1^ after 4 weeks of selection. The surviving shoots on SRM2+ medium were then individually separated and sub-cultured onto rooting medium RTM8 supplemented with cefotaxime (125 mg L^-1^) without kanamycin to initiate roots. Three separate transformation experiments were conducted with a total of 230 explants, where 80, 80 and 70 explants were used in experiment I, II and III, respectively.

### Screening of regenerated transgenic sweet orange plants

Leaf samples from regenerated sweet orange plants were immediately snap frozen in liquid nitrogen, freeze dried, and ground by TissueLyser II (Qiagen^®^). Genomic DNA (gDNA) was extracted using a DNeasy Plant Mini Kit (Qiagen^®^) following the manufacturer’s protocol. The extracted gDNA was used for PCR screening to identify transgenic sweet orange plants. gDNA from non-transgenic (NT) regenerated plants – induced from untransformed epicotyl explants – was used as a negative control. Primers (BEN0436F – 5’ CAAGCTCTTCAGCAATATCACG 3’ and BEN0437R – 5’ GACAGGTCGGTCTTGACAAA 3’) targeting the *NPTII* selectable marker gene were used in PCR reactions to amplify a ∼560 bp fragment from the left border portion of the empty CRISPR/Cas9 construct. The PCR reactions were performed using One*Taq*^®^ DNA Polymerase (NEB #M0480S) according to the manufacturer’s instructions and visualised on a 1% agarose gel electrophoresis. The transformation efficiency was calculated as the number of PCR-positive (presence of marker gene *NPTII*) regenerated plants divided by the total number of infected epicotyls with regenerated shoots, multiplied by 100.

### Statistical analysis

Both the shoot regeneration and rooting percentages were analysed using a one-way ANOVA followed by Tukey’s Honest Significant Difference (HSD) test in R Studio (version 2025.09.01+401). Post hoc groupings were generated, and Compact Letter Displays (cld) were used to indicate significance by lettering. Data with the same letter were not significantly different at *p* < 0.05.

## Results

### Factors enhancing regeneration competence of epicotyl explants

To obtain high-quality epicotyl explants, mature sweet orange seeds were placed on germination medium. Using fresh seed (≤6 months after fruit harvest), 91% of seeds began germinating within the first week, and germination was assessed finally at week four. We observed germination rates less than 50% when older sweet orange seeds (>6 months after fruit harvest) were used. Temperature during seed germination was found to be crucial; temperatures below 22°C delayed germination by 2-3 weeks, whereas 25 ± 1°C supported optimal germination, with dormancy broken in about one week. Four- to five-week-old light green seedlings were considered ideal sources of epicotyl explants for shoot regeneration (**Fig. 1a-c**). No shoots regenerated from slightly whitish explants (data not shown).

**Figure 1.**
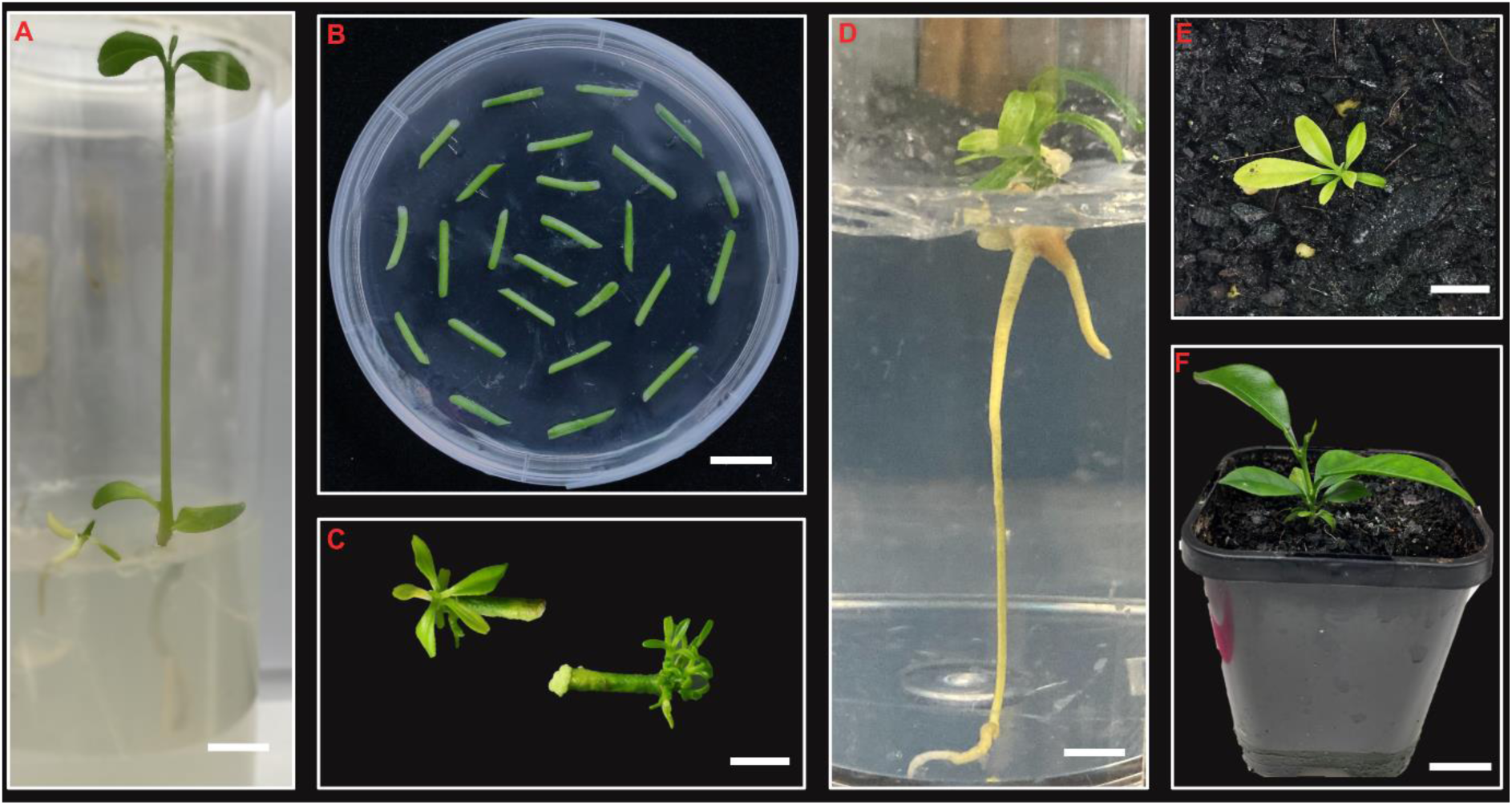
A schematic representation of sweet orange (*Citrus sinensis) in vitro* regeneration from epicotyl explants. (a) Optimal seedling growth for excising epicotyl explants, (b) dissected epicotyls (∼1 cm long) cultured in shoot regeneration media, (c) regenerated shoots from the cut end of the epicotyls, (d) elongated *in vitro* roots from regenerated shoots, (e) acclimatisation of plantlets 5 days after transfer to soil, (f) fully grown healthy sweet orange plantlet 8 weeks after transfer to soil. Scale bar = 1 cm.

### Determining the best-performing plant growth regulators (PGRs) for shoot regeneration

Previous research has shown that PGRs such as cytokinins and auxins are effective for inducing shoots from citrus explants. To determine the best-performing PGRs, the frequency of shoot regeneration from epicotyl explants was evaluated on SRMs containing varying concentrations and combinations of BAP, IAA, and 2,4-D (**Table 1**). Shoot initiation began about 7 days after culture (**Fig. 1c**), and final counts were taken at 6 weeks. Newly regenerated shoots exhibited several patterns, including emergence from one or both sides of the epicotyl cut edges, formation as single or clustered shoots, and in a few cases, we observed callus developed at one end of the epicotyl while shoots originated from the opposite end (**Supplementary Fig. S2**). The highest shoot regeneration (90.6 ± 1.73%) was observed for SRM2 supplemented with 3.0 mg L^-1^ BAP, which is significantly higher than the other media tested (**Table 1**). Reducing the concentration of BAP to 2.0 mg L^-1^, with (SRM3) or without (SRM1) the addition of IAA (0.5 mg L^-1^), lowered the regeneration rate, while BAP in combination with 2,4-D (SRM4) failed to regenerate any shoots but instead only formed callus (**Table 1**). Explants cultured on SRM3 (2.0 mg L^-1^ BAP + 0.5 mg L^-1^ IAA) generated both shoots and callus. These findings indicate that cytokinin alone, in the form of BAP and at an optimal concentration, is effective for shoot regeneration in sweet orange and the auxin-cytokinin interactions vary depending on the type of auxin used.

**Table 1.** *In vitro* shoot regeneration of sweet orange (*Citrus sinensis)* epicotyl explants cultured on various shoot regeneration media (SRM) with different combinations of plant growth regulators (PGRs).

| Media | Concentrations of PGRs ($\text{mg L}^{-1}$) | | | Shoot regeneration rate (%) |
| --- | --- | --- | --- | --- |
|  | Cytokinin (BAP) | Auxin (IAA) | Auxin (2,4-D) |  |
| SRM1 | 2.0 | 0 | 0 | $75.0 \pm 1.73^b$ |
| SRM2 | 3.0 | 0 | 0 | $90.6 \pm 1.73^a$ |
| SRM3 | 2.0 | 0.5 | 0 | $77.1 \pm 1.0^b$ |
| SRM4 | 2.0 | 0 | 0.25 | $0 \pm 0^c$ |

Values are presented as means ± SE. Different superscript letters indicate statistically significant differences between SRMs (One-way ANOVA followed by Tukey's HSD test, $p < 0.05$ ). $n = 3$ (of 32 explants of each replicates).

After 2-3 weeks of culture, regenerated shoots were detached from the explants and transferred to fresh SRM of the same composition for further elongation. The number of shoots regenerated per epicotyl varied widely, ranging from a single shoot to as many as 17 shoots (**Supplementary Fig. S2**). We observed that clustered shoots were generally weaker, and many failed to continue growing when detached as a cluster from the epicotyl; typically, only two to three centrally positioned shoots within a cluster survived and developed leaves. To improve recovery of healthier shoots, individual shoots were separated from clusters and sub-cultured on fresh media, and we observed survival of a much higher number of shoots (data not shown). For subsequent rooting, shoots approximately 1-2 cm long were carefully excised and transferrd onto RTM media (**Fig. 1d**).

### Optimisation of PGR ratios for robust root formation

Different strengths of basal rooting media without PGRs were initially tested for rooting the *in vitro* shoots. A significantly higher rooting rate (29.2 ± 0.95%) was observed in shoots cultured on half-strength MT medium compared with those cultured on full-strength MT medium (19 ± 1.91%) (**Supplementary Table S1**). Therefore, half-strength MT medium was selected as the basal medium for all further RTMs used in this study.

Root induction began 2 weeks after transferring *in vitro* shoots to the various RTMs, containing different concentrations and combinations of IBA and NAA (**Table 2**). Rooting occurred in all RTMs tested, but rooting percentages varied significantly among media. IBA alone at lower concentrations (0.5 - 1.0 mg L^-1^) induced roots in 33-42% of shoots, whereas 2.0 mg L^-1^ IBA yielded only 13% rooting. In comparison, NAA-only RTMs generally produced lower rooting percentages (8 - 29%).

**Table 2.**
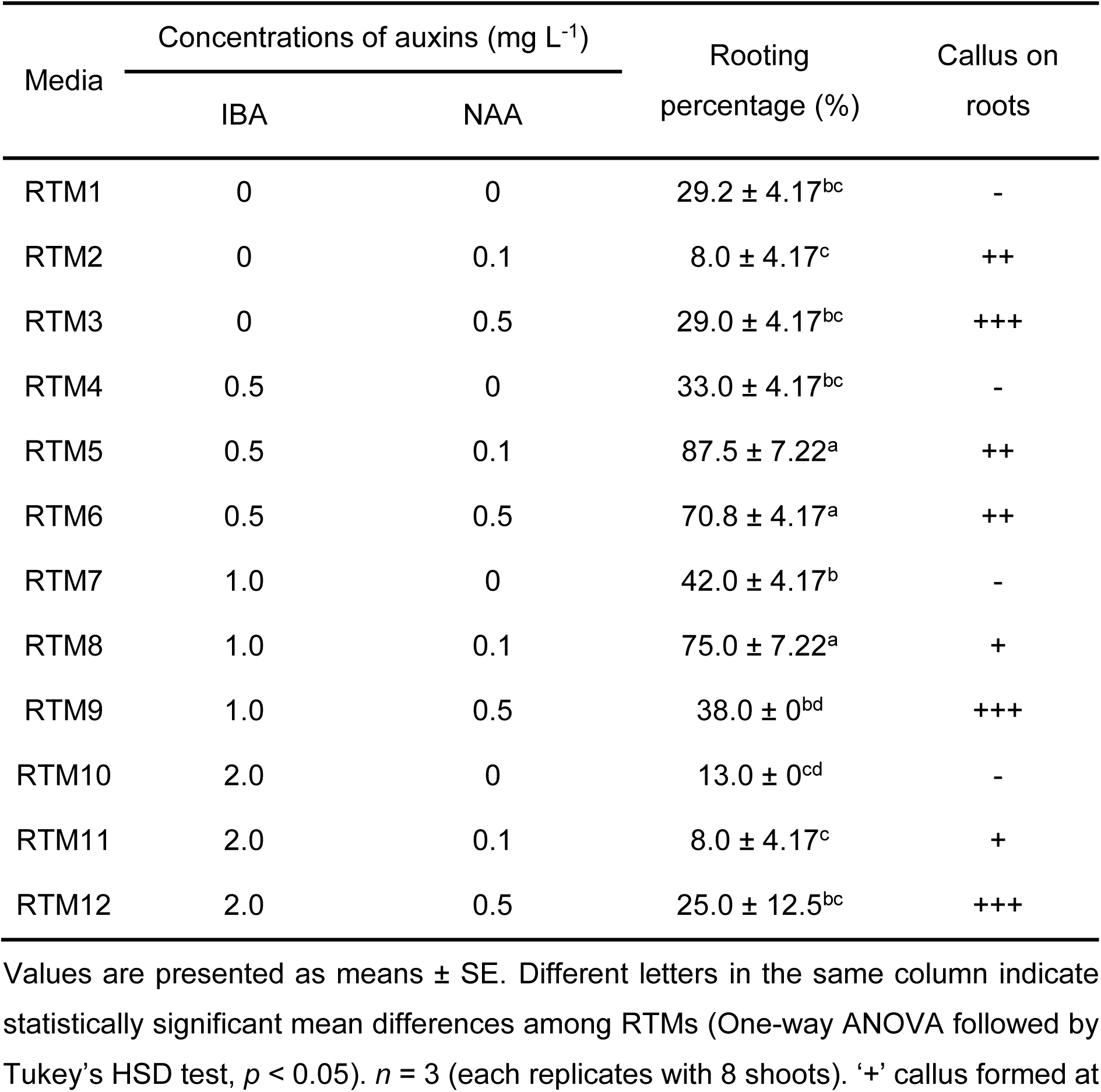
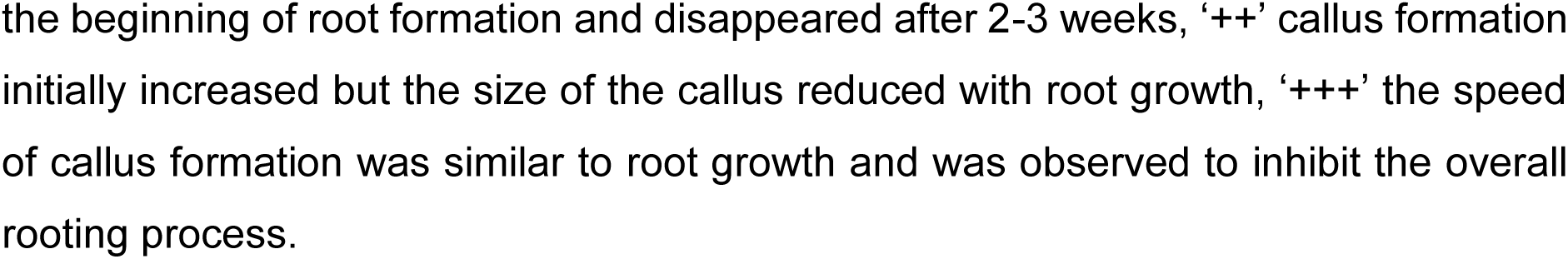
Rooting responses of regenerated sweet orange (*Citrus sinensis*) shoots cultured on different rooting media (RTM) containing a range of concentrations of the auxins, indole-3-butyric acid (IBA) and 1-naphthalene acetic acid (NAA).

| Media | Concentrations of auxins (mg L<sup>-1</sup>) |  | Rooting percentage (%) | Callus on roots |
| --- | --- | --- | --- | --- |
|  | IBA | NAA |  |  |
| RTM1 | 0 | 0 | $29.2 \pm 4.17^{bc}$ | - |
| RTM2 | 0 | 0.1 | $8.0 \pm 4.17^c$ | ++ |
| RTM3 | 0 | 0.5 | $29.0 \pm 4.17^{bc}$ | +++ |
| RTM4 | 0.5 | 0 | $33.0 \pm 4.17^{bc}$ | - |
| RTM5 | 0.5 | 0.1 | $87.5 \pm 7.22^a$ | ++ |
| RTM6 | 0.5 | 0.5 | $70.8 \pm 4.17^a$ | ++ |
| RTM7 | 1.0 | 0 | $42.0 \pm 4.17^b$ | - |
| RTM8 | 1.0 | 0.1 | $75.0 \pm 7.22^a$ | + |
| RTM9 | 1.0 | 0.5 | $38.0 \pm 0^{bd}$ | +++ |
| RTM10 | 2.0 | 0 | $13.0 \pm 0^{cd}$ | - |
| RTM11 | 2.0 | 0.1 | $8.0 \pm 4.17^c$ | + |
| RTM12 | 2.0 | 0.5 | $25.0 \pm 12.5^{bc}$ | +++ |

Values are presented as means ± SE. Different letters in the same column indicate statistically significant mean differences among RTMs (One-way ANOVA followed by Tukey's HSD test, $p < 0.05$ ). $n = 3$ (each replicates with 8 shoots). '+' callus formed at
the beginning of root formation and disappeared after 2-3 weeks, ‘++’ callus formation initially increased but the size of the callus reduced with root growth, ‘+++’ the speed of callus formation was similar to root growth and was observed to inhibit the overall rooting process.

When evaluating the effects of combined IBA and NAA on rooting percentages, a similar trend was observed where IBA in lower concentration performed better than in higher concentration, with the addition of NAA generally increasing the rooting percentages. Significantly higher rooting percentages (70.8 ± 4.17% to 87.5 ± 7.22%) were obtained in RTM5, RTM6 and RTM8, which contained IBA at 0.5 or 1.0 mg L^-1^ and NAA at 0.1 or 0.5 mg L^-1^, when compared to all other RTMs. No significant differences were detected among these three RTMs. Overall, while roots formed across media, optimised auxin concentrations and combinations markedly increased rooting frequency.

Consistent with the effects of cytokinin concentration and auxin-cytokinin interactions on shoot regeneration, auxin concentration, type, and combination significantly affected root growth and morphology. While we mostly observed development of a single tap root from regenerated shoots, diverse root morphologies were observed across treatments with IBA and NAA, varying with both their combinations and concentrations (**Fig. 2**). Although NAA, in combination with IBA, increased the number of shoots inducing roots, the presence of NAA also promoted callus formation on the roots. At high concentrations of NAA (0.5 mg L^-1^) the rate of callus formation significantly affected root elongation (**Fig. 2a**), while lower concentration of NAA (0.1 mg L^-1^) enabled roots to elongate to 5-6 cm within 4 weeks without significant callus formation (**Fig. 2d**). Based on our continuous observation, RTM8 (1.0 mg L^-1^ IBA + 0.1 mg L^-1^ NAA) was selected as the best-performing rooting medium because it produced a consistently higher rooting response (75.0 ± 7.22%), but with longer and more vigorous roots than RTM5 (0.5 mg L^-1^ IBA + 0.1 mg L^-1^ NAA) and RTM6 (0.5 mg L^-1^ IBA and 0.5 mg L^-1^ NAA).

**Figure 2.**
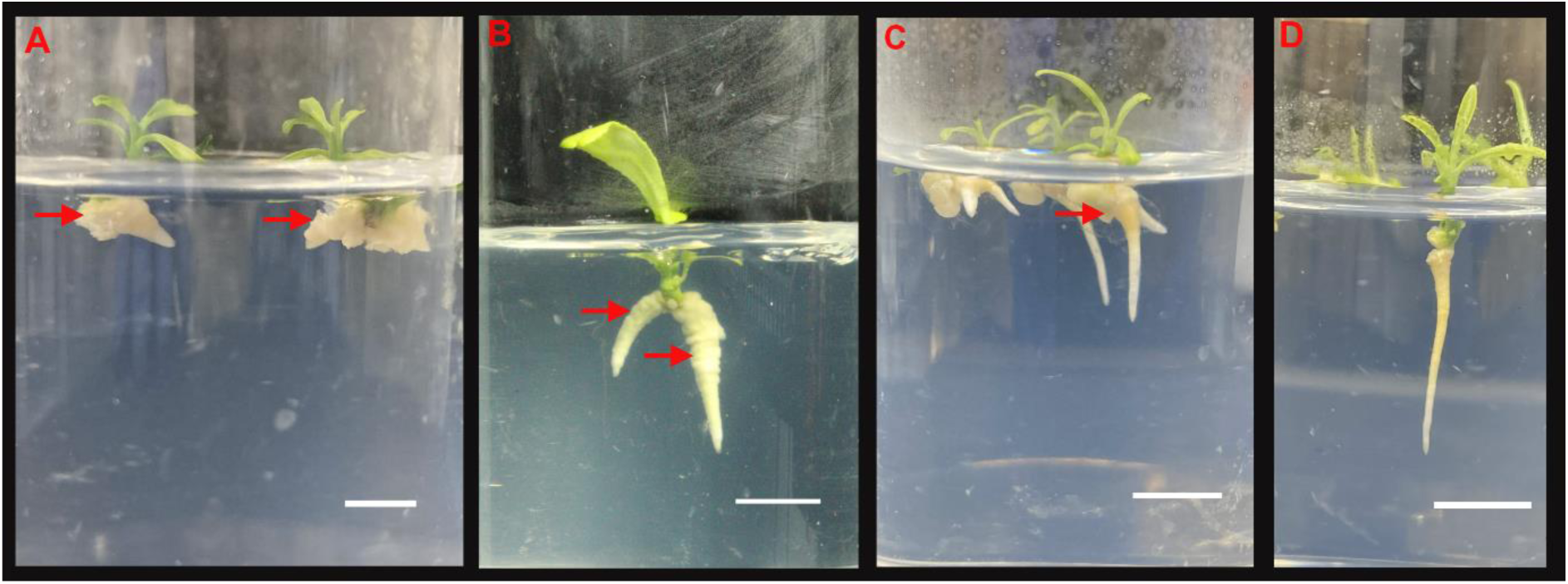
Diverse root morphologies of epicotyl-regenerated sweet orange (*Citrus sinensis*) shoots cultured on different rooting media. (a) Callus formation at the basal end of the shoot with no visible root development in rooting media mostly containing 0.5 mg L-^1^ NAA, (b) root formation observed, however, excessive callus formation reduced shoot vigour, (c) callus present at the basal region during early rooting, followed by elongation of roots and gradual disappearance of callus tissue, (d) absence of callus formation with well-developed, elongated roots. (a-c) Callus tissues are indicated by red arrows. Scale bar = 1 cm.

### Efficient regeneration of transgenic sweet orange plants using the optimised regeneration protocol

We evaluated the efficiency of our best-performing SRM and RTM in regenerating sweet orange plants from *Agrobacterium*-transformed epicotyl explants. Transgenic sweet orange plants were successfully recovered following transformation of sweet orange epicotyl explants with an *Agrobacterium* suspension harbouring an empty CRISPR/Cas9 construct (**Table 3** and **Fig. 3a-e**). The regeneration process was comparable to that of untransformed explants; however, the inclusion of selection pressure to facilitate the recovery of transgenic shoots impacted on the shoot regeneration rate. The shoot regeneration rate reached 41.5% on selection medium SRM2+ when *Agrobacterium*-treated epicotyls were selected with 100 mg L^-1^ kanamycin (**Table 3**). Pre-trial results indicated that 50 mg L^-1^ kanamycin did not provide sufficient selection pressure, while fewer than 5% of explants formed shoots on 150 mg L^-1^ kanamycin.

**Figure 3.**
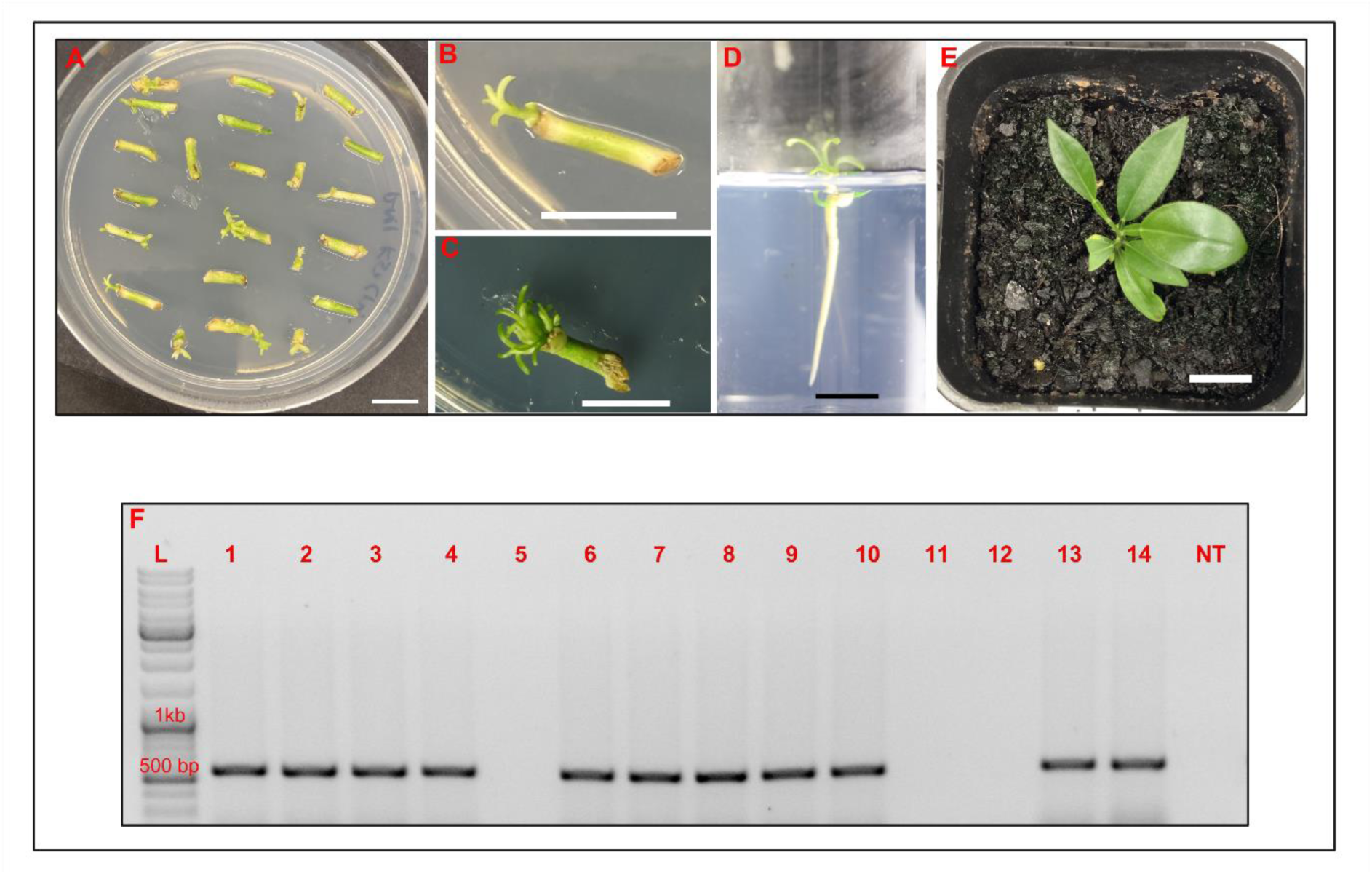
Recovery and molecular confirmation of transgenic sweet orange (*Citrus sinensis*) lines. (a) Epicotyl explants infected with the empty CRISPR/Cas9 construct cultured on regeneration media under selection, (b-c) regenerated transgenic shoots on SRM2+ selection medium, (d) rooted transgenic plantlets, (e) acclimatised transgenic plantlets 8 weeks after transfer to soil, (f) PCR confirmation of *Cas9* integration in regenerated sweet orange plants. Lane left to right; L = gene ruler ladder mix indicated with 1 kb and 500 bp bands, transgenic lines are labelled 1-14, and NT indicates a non-transformed control. *n* = 230 (total number of explants used in three experiments). (a-e) Scale bar = 1 cm.

**Table 3.** Efficiency of best-performing shoot regeneration (SRM2+) and rooting (RTM8) media for recovering transgenic sweet orange (*Citrus sinensis*) plants.

| Treatment | Shoot regeneration rate (%) | Rooting percentage (%) | Transformation efficiency |
| --- | --- | --- | --- |
| Control | 87.0 ± 0.67 | 73.0 ± 2.1 | N/A |
| Empty CRISPR/Cas9 construct | 41.5 ± 0.06 | 69.0 ± 1.1 | 4.8 ± 0.2% |

Values are presented as means ± SE. Transformation efficiency (TE) = (positive regenerated shoots confirmed by PCR / the total number of treated epicotyl explants (n)) × 100. n = 230 (3 independent experiments, totaling 230 explants across all trials).

The regenerated shoots obtained from selection medium successfully developed roots when manually excised and cultured on the high-performing rooting medium, RTM8 (**Fig. 3c**). Cefotaxime, which was included in RTM8 to suppress *Agrobacterium* overgrowth, did not adversely affect rooting of transformed (selection-derived) shoots when compared with non-transformed (control) shoots (**Table 3**). When cultured on RTM8, 69% of the transformed shoots developed roots, demonstrating the effectiveness of this RTM for the recovery of *Agrobacterium*-treated epicotyls.

A total of 14 plants were tested to confirm transformation, obtained from the three experiments, 5 from experiments I and II, and 4 from experiment III. gDNA from all these 14 plants was isolated from the leaf tissue of regenerated sweet orange plants and the *NPTII* gene, providing kanamycin resistance, was amplified by PCR. Eleven out of fourteen regenerated plantlets produced the expected ∼560 bp amplicon corresponding to the *NPTII* gene (**Fig. 3f**), confirming successful T-DNA integration. We found a transformation efficiency of 4.8% as the number of PCR-positive events relative to the total number of treated epicotyl explants (**Table 3**).

### Acclimatisation of regenerated plantlets

*In vitro* regenerated sweet orange plantlets, both the transformed with empty CRISPR/Cas9 construct (**Fig. 3a-e**) and the non-transformed (**Fig. 1**), were transferred to ex vitro condition to complete the acclimatisation process. Healthy rooted plantlets, 20 and 16 plantlets of each of the non-transformed and transformed explant-derived regenerated plants, respectively, were acclimatised after removal from the rooting medium and transplanting into autoclaved potting mixture. We observed in a pre-experiment trial that excessive watering resulted in root damage, whereas maintaining a slightly drier substrate during the first few days promoted better establishment. After 4 weeks, approximately 90% of regenerated sweet orange plantlets survived and continued normal growth and development (**Fig. 1f** and **Fig. 3e**).

## Discussion

A wide range of explants such as epicotyls, hypocotyls, stems, leaves, and roots have previously been used for *in vitro* regeneration of commercial citrus species, although epicotyls are consistently reported as one of the most responsive and reliable explant sources ^[8,15]^. However, *in vitro* seed germination plays an important role in supplying competent explants, which is affected by seed age, seed treatment, and germination culture conditions ^[8,37]^. Previous research has shown that the developmental stage of sweet orange seedlings affects subsequent regeneration efficiency from epicotyl explants ^[25,38]^.

In this study, we applied optimum germination and culture practices to obtain high-quality epicotyl explants and evaluated their regeneration responses under different PGR treatments. The same explant system was subsequently used to generate transgenic sweet orange plants.

### Cytokinin-induced shoot regeneration in sweet orange

Citrus regeneration is strongly influenced by PGRs, with genotype-dependent responses often complicating protocol optimisation. Among cytokinin, BAP is widely recognised as the most effective for inducing shoots in citrus, whereas kinetin and zeatin typically yield lower regeneration rates ^[39]^. For this reason, we trialled SRMs containing mainly BAP to induce shoots from sweet orange epicotyl explants and found significantly higher shoot induction (90.6 ± 1.73%) at high concentrations of BAP (3.0 mg L^-1^). However, prior reports indicate that high BAP levels (≥3.0 mg L^-1^) can be phytotoxic to explant growth resulting in reduced shoot proliferation and mean shoot number in several *Citrus* species ^[40,41]^. In contrast, Jardak et al ^[42]^ reports higher shoot regeneration of ‘Maltese half-blood’ (*C. sinensis*) epicotyls with BAP at 2.0 mg L^-1^ and 4.0 mg L^-1^. We did not observe shoot reduction or any toxicity in high BAP (3.0 mg L^-1^) here in sweet orange highlighting the species-specific and cultivar differences common in citrus tissue culture.

Combinations of BAP with auxins such as IAA or NAA have improved shoot formation in some citrus species ^[15,39]^, but in other citrus species BAP in combination with the auxins NAA or 2,4-D promote callus formation ^[43,44]^. In this study we observed both shoot regeneration and callus formation from sweet orange epicotyls on BAP+IAA supplemented-media, and only callus formation on media containing BAP+2,4-D. Therefore, our results indicate that BAP alone is sufficient to stimulate shoot organogenesis in sweet orange explants, while the addition of auxins induces callus formation and does not enhance shoot regeneration. This is likely due to adequate endogenous auxin levels in sweet orange ^[45]^ and the central role of cytokinins in cell division, differentiation, and nutrient-stress signalling pathways ^[46,47]^. Therefore, a simplified BAP-based medium offers an efficient and practical approach for shoot regeneration in sweet orange.

### Optimisation of auxin-mediated rooting in sweet orange

After successful shoot regeneration, *in vitro* rooting is often a major bottleneck for many crop species ^[48,49]^, particularly perennial woody species such as *Citrus* ^[50]^. Achieving reliable rooting is therefore essential for completing the regeneration cycle and improving the efficiency of both commercial propagation and transgenic research. In this study, we demonstrated an optimised auxin-based rooting strategy that significantly improved root induction in recalcitrant sweet orange.

Historically, rooting responses in citrus have been inconsistent, with several species–including mandarin, sour orange, and grapefruit–showing failure to initiate roots, or producing roots with low survival during acclimatisation ^[51–53]^. In contrast, we show that sweet orange exhibits enhanced rooting when regenerated shoots are transferred to reduced-strength basal media supplemented with appropriate auxins combinations. Among the auxins tested, IBA consistently promoted higher rooting efficiency compared with NAA when applied individually, supporting earlier evidence that IBA is more effective in inducing roots in citrus ^[41]^. A synergistic effect was observed when IBA and NAA were combined; however, their concentrations were critical for optimal outcome. Lower concentrations of IBA (0.5 mg L^-1^) combined with NAA (0.1-0.5 mg L^-1^) increased rooting percentage but simultaneously caused callus formation and restricted root elongation. This pattern aligns with the known auxin signalling dynamics, where excessive exogenous auxin can disrupt normal organogenic patterning and promote callus formation by triggering cell dedifferentiation and reprogramming ^[54,55]^. Conversely, increasing IBA levels to 1.0 mg L^-1^ while lowering NAA to 0.1 mg L^-1^ produced the highest rooting rates and the longest roots while minimising callus formation highlighting the dose-dependent nature of PGR interactions in citrus. This agrees with previous reports showing auxin-specific and genotype-dependent rooting responses in citrus species ^[11,40,56]^. For example, different concentrations of IAA significantly affect rooting of alemow (*Citrus macrophylla*) shoots, but IBA alone enhances rooting percentages of ‘Cleopatra’ mandarin (*Citrus reshni*) shoots ^[40]^.

Although NAA facilitated root initiation, higher concentrations negatively affected root architecture by promoting callus at the shoot base and suppressing root growth, consistent with findings in other plant species ^[48,57]^. In contrast, IBA provides a more sustained and physiologically regulated auxin supply that has been associated with enhanced adventitious root development *in vitro* ^[58]^. Overall, our results demonstrate that precise optimisation of IBA-NAA combinations can overcome *in vitro* rooting challenges in sweet orange and substantially streamline the regeneration pipeline for applications like genetic transformation.

### Recovery of transgenic sweet orange plants using the optimised regeneration system

Genome editing and genetic transformation research in many crops, including citrus, is limited by inefficient regeneration protocols and the difficulty of recovering stable transgenic plants ^[59]^. In citrus, factors such as regeneration medium, tissue culture conditions, and explant type are known to strongly influence transformation sources ^[33,60,61]^. In this study, we evaluated the ability of high-performing shoot regeneration and rooting media to improve recovery of transgenic sweet orange plants. Best-performing PGR combinations were first identified using sweet orange epicotyl explants and then applied to *Agrobacterium*-mediated epicotyl transformation experiments.

Shoots regenerated from explants transformed with an empty CRISPR/Cas9 construct exhibited healthy rooting responses on the optimised auxin-supplemented rooting medium, comparable to non-transformed controls. Using this improved system, transgenic sweet orange plants were successfully recovered within 3-4 months after transformation, demonstrating that the refined rooting strategy accelerates the regeneration pipeline. The transformation efficiency of 4.8% obtained in this study falls within the range previously reported for citrus, which varies widely (1-55%) depending on explant type, cultivar, bacterial strain, and regeneration conditions ^[13,17,62]^. Furthermore, the successful application of an empty CRISPR/Cas9 construct confirms the suitability of this regeneration system for future genome-editing work in citrus.

## Conclusions

In summary, this study establishes an efficient *in vitro* regeneration system capable of producing whole sweet orange plants from both *Agrobacterium*-infected and non-infected epicotyl explants of sweet orange, a commercially important citrus variety. For shoot induction, media supplemented with BAP alone proved sufficient to stimulate reliable regeneration. In contrast, successful root formation required precise optimisation of auxin levels, with auxin-rich rooting media significantly enhancing root initiation and development. The best-performing media also enabled recovery of transgenic plants within a comparatively short timeframe, demonstrating its suitability for genetic transformation and future genome-editing applications in citrus.

## Author contributions

Study concept and design: Datta J, Kerr SC; Data collection, interpretation, analysis and visualisation: Datta J; supervision: Bhowmik SD, Williams B, Kerr SC; draft manuscript preparation: Datta J; resources and validation: Bhowmik SD, Williams B; project administration and funding acquisition: Kerr SC. All authors reviewed the results and approved the final version of the manuscript.

## Acknowledgements

The authors acknowledge financial support for this research from Queensland University of Technology (QUT) Postgraduate Research Award (QUTPRA), National Tree Genomics (AS17000) project, and Genetics for Next Generation Orchards (AS23003) project which is funded through Hort Innovation Frontiers with co-investment from Queensland University of Technology, University of Queensland, the Queensland Alliance for Agriculture and Food Innovation (QAAFI), Western Sydney University, Murdoch University, Adelaide University, and The University of Western Australia; Department of Primary Industries (Queensland), Department of Primary Industries and Regional Development (Western Australia), and Northern Territory Department of Agriculture and Fisheries; Bioeconomy Science Institute, CIRAD, AbacusBio; and contributions from the Australian Government. The authors also thank Mr. Malcolm Smith (Bundaberg Research Station, Department of Primary Industries, Queensland Government) for providing the plant material used in tissue culture experiments, and Ms. Sally Roden (Centre for Agriculture and the Bioeconomy, Queensland University of Technology) for providing the plasmid of pCAMBIA2300 binary vector.

## Data availability

All data generated or analysed during this study are included in this published article and its supplementary information files.

