## Supplementary materials for "An *in vitro* regeneration system with efficient rooting in sweet orange (*Citrus sinensis*) supports recovery of transgenic plants"

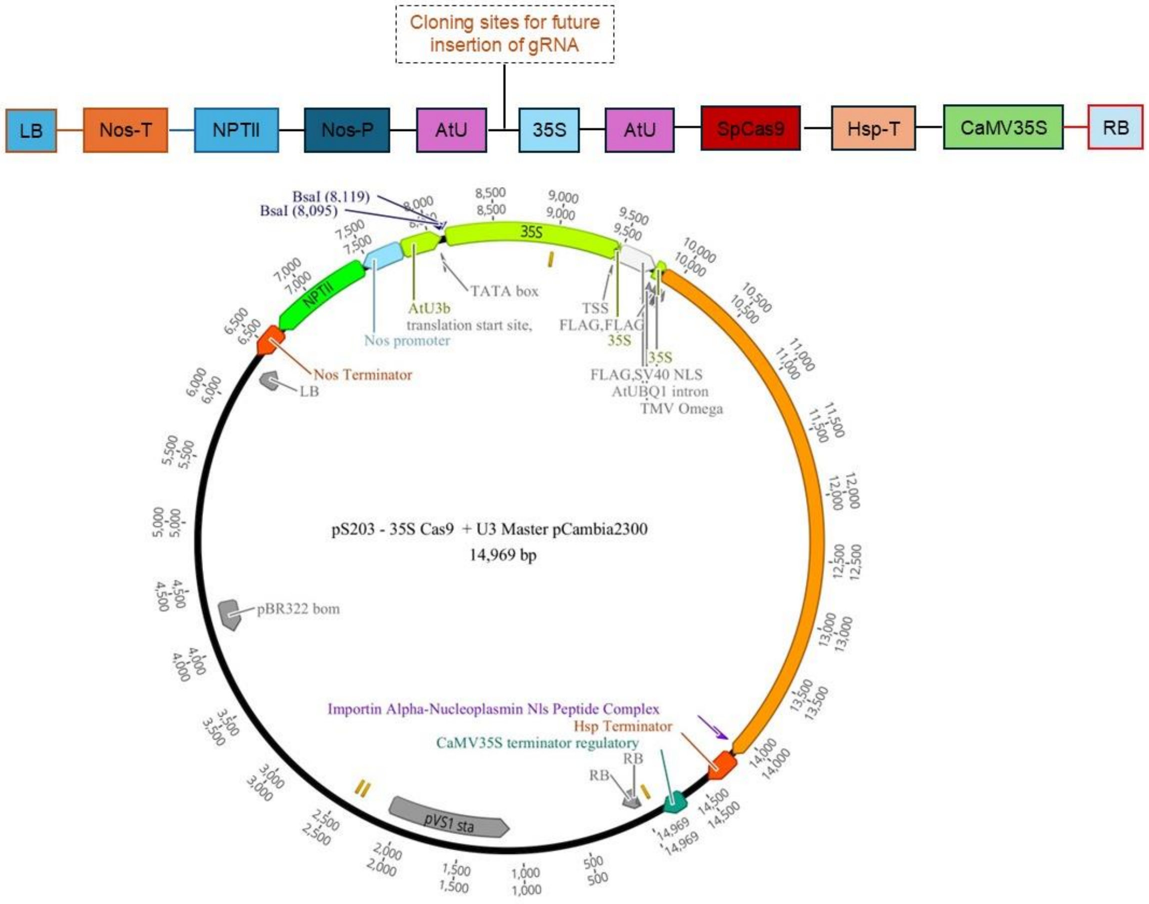


**Supplementary Figure S1**. Schematic presentation of the plasmid map of the empty CRISPR/Cas9 construct. pS203 (modified pCAMBIA2300) containing Cas9 expression cassette driven by the CaMV 35S promoter. Cloning sites are located between the AtU and CaMV 35S promoters for future integration of single or multiple gRNA. RB: right border.

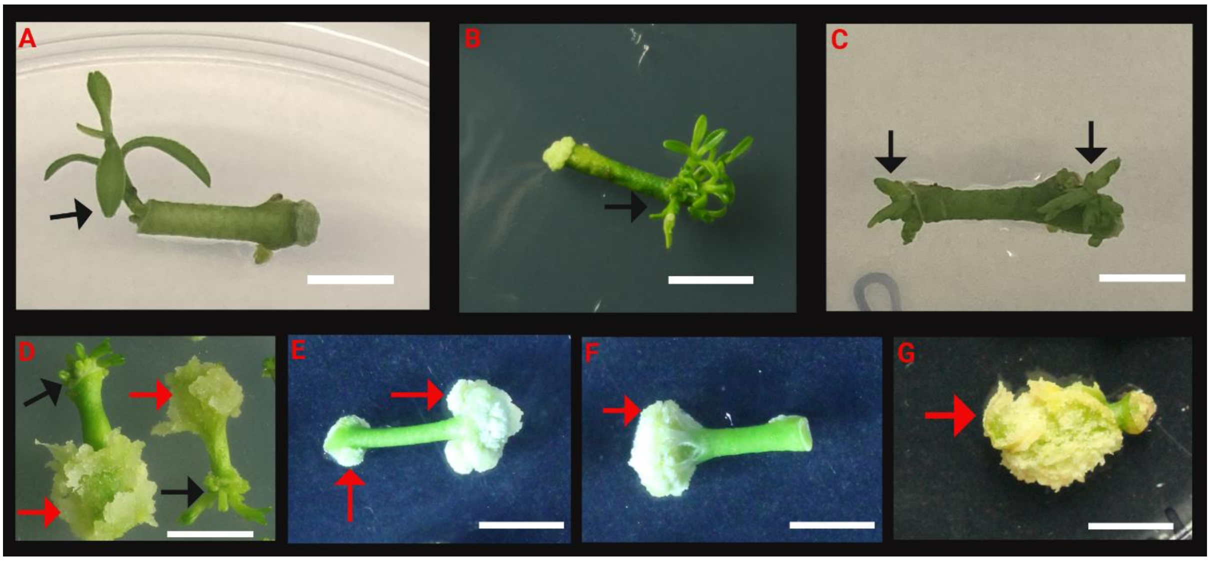


**Supplementary Figure S2.** Different patterns of shoot regeneration from sweet orange epicotyl explants cultured on various shoot regeneration media. (a) Single shoot induction from an epicotyl explant, (b) clustered shoot formation; (c) shoot formation from both ends of an epicotyl explant, (d) callus formation at one end with shoot induction at the opposite end, (e-g) callus formation without shoot regeneration: (e) callus formed at both ends, (f) callus formed at one end, and (g) callus covering the entire epicotyl explant. Black arrows indicate regenerated shoots, while red arrows indicate callus. Scale bar = 0.5 cm.

**Supplementary Table S1.** Half-strength MT medium shows increased rooting rate over full-strength MT medium.

| MT strength | IBA | NAA | Rooting (%) |
|---|---|---|---|
| Full | 0 | 0 | $19.0 \pm 1.9^{b}$ |
| Half | 0 | 0 | $29.2 \pm 0.95^{a}$ |

Values are presented as mean ± SE. Different superscript letters in the same column denote statistically significant differences between MT media (One way ANOVA, $p < 0.05$). $n = 3$ (of 8 regenerated shoots per replicates).
